# Evolution of genes neighborhood within reconciled phylogenies: an ensemble approach

**DOI:** 10.1101/026310

**Authors:** Cedric Chauve, Yann Ponty, João Paulo Peirera Zanetti

## Abstract

**Context:** The reconstruction of evolutionary scenarios for whole genomes in terms of genome rearrangements is a fundamental problem in evolutionary and comparative genomics. The DeCo algorithm, recently introduced by Bérard *et al.*, computes parsimonious evolutionary scenarios for gene adjacencies, from pairs of reconciled gene trees. However, as for many combinatorial optimization algorithms, there can exist many co-optimal, or slightly sub-optimal, evolutionary scenarios that deserve to be considered.

**Contribution:** We extend the DeCo algorithm to sample evolutionary scenarios from the whole solution space under the Boltzmann distribution, and also to compute Boltzmann probabilities for specific ancestral adjacencies.

**Results:** We apply our algorithms to a dataset of mammalian gene trees and adjacencies, and observe a significant reduction of the number of syntenic conflicts observed in the resulting ancestral gene adjacencies.

## Background

The reconstruction of the evolutionary history of genomic characters along a given species tree is a long-standing problem in computational biology. This problem has been well studied for several types of genomic characters, for which efficient algorithms exist to compute parsimonious evolutionary scenarios; classical examples include genes and genomes sequences [1], gene content [2], and gene family evolution [3, 4]. Recently, Bérard *et al.* [5] extended the corpus of such results to syntenic characters. They introduced the notion of adjacency forest, that models the evolution of gene adjacencies within a phylogeny, motivated by the reconstruction of the architecture of ancestral genomes, and described an efficient dynamic programming (DP) algorithm, called `DeCo`, to compute parsimonious adjacency evolutionary histories. So far, `DeCo` is the only existing tractable model that considers the evolution of gene adjacencies within a general phylogenetic framework: other tractable models of genome rearrangements accounting for a given species phylogeny are either limited to single-copy genes and ignore gene-specific events [6], assume restrictions on the gene duplication events, such as considering only whole-genome duplication (see [7] and references there), or require a dated species phylogeny [8].

From a methodological point of view, most existing algorithms to reconstruct evolutionary scenarios along a species tree in a parsimony framework rely on dynamic-programming along this tree, whose introduction can be traced back to Sankoff in the 1970s (see [9] for a recent retrospective on this topic). Recently, several works considered more general approaches for such parsimony problems that either explore a wider range of values for combinatorial parameters of parsimonious models [10] or consider several alternate histories for a given instance, chosen for example from the set of all possible co-optimal scenarios or from the whole solution space, including suboptimal solutions (see [11, 12, 13] for examples of this approach for the gene tree/species tree reconciliation problem).

The present work follows the later approach and extends the `DeCo` DP scheme toward an exploration of the whole solution space of adjacency histories, under the Boltzmann probability distribution, that assigns a probability to each solution defined in terms of its parsimony score. This principle of exploring the solution space of a combinatorial optimization problem under the Boltzmann probability distribution is sometimes known as the “Boltzmann ensemble approach”. It was initially introduced in the context of RNA folding, where the probability of any given conformation at the thermodynamic equilibrium follows a Boltzmann distribution, i.e. a conformation *s* is observed for a given RNA *w* with probability 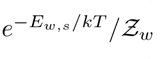 where *E*_*w,s*_ is the free-energy of conformation *s* over *w*, *k* is the Boltzmann constant, *T* is the temperature, and *Z*_*w*_ is the partition function of *w*. This latter quantity can be seen as a renormalization factor, and is key in the study of RNA thermodynamics, but its computation involves summing over an exponential number of conformations compatible with the RNA sequence. A major paradigm shift occurred in RNA research when McCaskill [14] showed in 1990 how an efficient algorithm for the partition function could be adapted from a DP scheme for energy minimization through a simple change of algebra. This seminal work also introduced a variant of the *inside-outside* algorithm [15] for computing base-pairing probabilities.

While this Boltzmann ensemble approach has been used for a long time in RNA structure analysis, to the best of our knowledge it is not the case in comparative genomics, where exact probabilistic models have been favoured recently [16, 17]. However, probabilistic models still pose computational challenges for large datasets, and so far a probabilistic model does not exist for gene adjacencies, which motivates our work. In the specific case of the `DeCo` model, the ability to explore alternative co-optimal or slightly sub-optimal solutions is crucial. Indeed, as `DeCo` models gene adjacencies, each ancestral gene can only be adjacent to at most two other genes, which is not considered in `DeCo`. However, the initial experiments using `DeCo` on mammalian gene trees resulted in hundreds of ancestral genes were involved in more than two ancestral gene adjacencies [5]. This raises the question of filtering inferred ancestral adjacencies to reduce the level of syntenic conflict, which can be done on the basis of their Boltzmann probabilities. We reason that some of the erroneously-predicted adjacencies may result from combinatorial optimization artifacts and that features of a gene adjacency parsimonious evolutionary scenario that are not robust to considering alternative equivalent, or slightly worse, solutions should be considered as dubious.

## Methods

### Models

A *phylogeny* is a rooted tree which represents the evolutionary relationships of a set of elements represented by its nodes: internal nodes are ancestors, leaves are extant elements, and edges represent direct descents between parents and children. We consider here three kinds of phylogenies (illustrated in Figure 1): species trees, reconciled gene trees and adjacencies trees/forests. Trees we consider are always rooted. For a tree *T* and a node *x* of *T*, we denote by *T* (*x*) the subtree rooted at *x*. If *x* is an internal node, we assume it has either one child, denoted by *a*_*x*_, or two children, denoted by *a*_*x*_ and *b*_*x*_.

**Figure 1.**
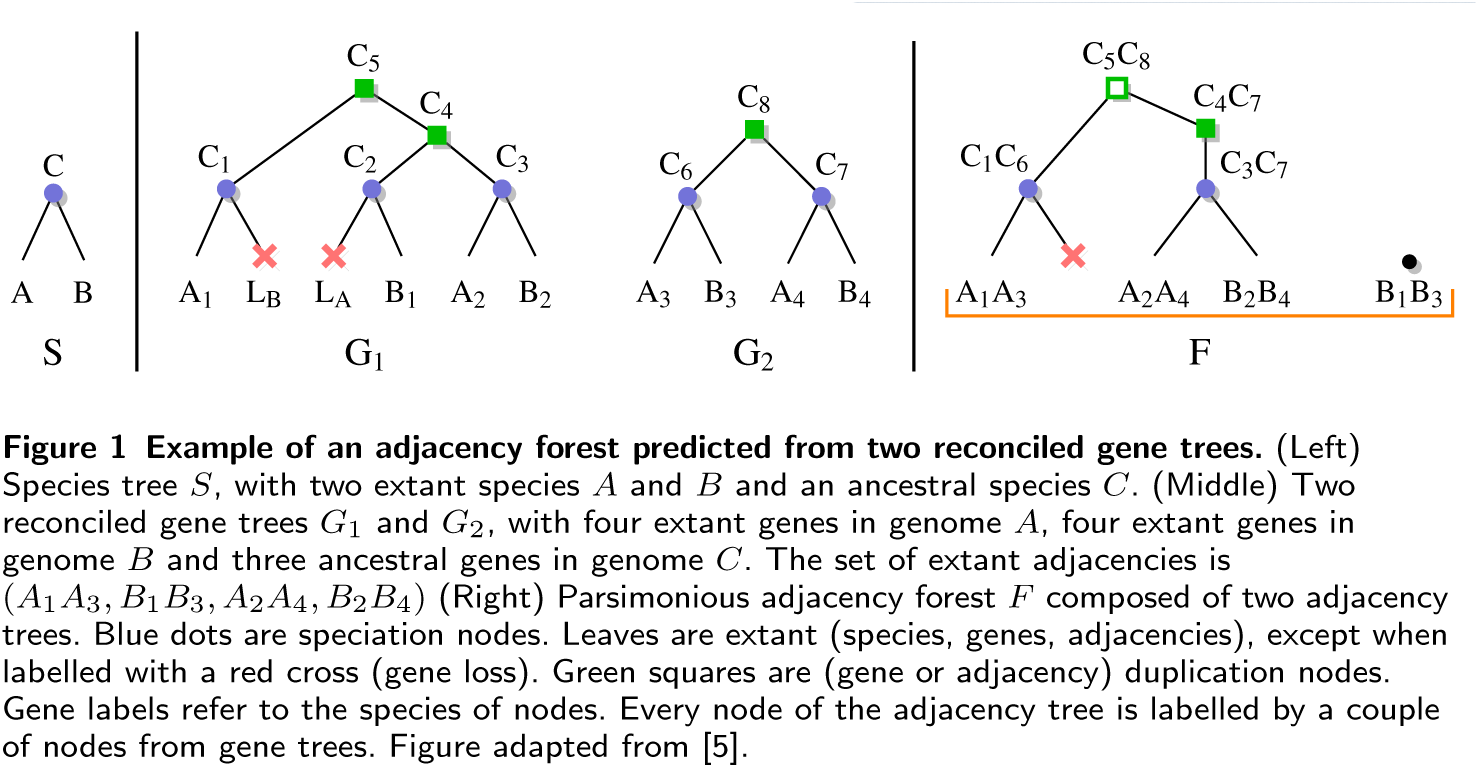
Example of an adjacency forest predicted from two reconciled gene trees. (Left) Species tree *S*, with two extant species *A* and *B* and an ancestral species *C*. (Middle) Two reconciled gene trees *G*_1_ and *G*_2_, with four extant genes in genome *A*, four extant genes in genome *B* and three ancestral genes in genome *C*. The set of extant adjacencies is (*A*_1_*A*_3_; *B*_1_*B*_3_; *A*_2_*A*_4_; *B*_2_*B*_4_) (Right) Parsimonious adjacency forest *F* composed of two adjacency trees. Blue dots are speciation nodes. Leaves are extant (species, genes, adjacencies), except when labelled with a red cross (gene loss). Green squares are (gene or adjacency) duplication nodes. Gene labels refer to the species of nodes. Every node of the adjacency tree is labelled by a couple of nodes from gene trees. Figure adapted from [5].

#### Species trees

A species tree *S* is a binary tree that describes the evolution of a set of related species, from a common ancestor (the root of the tree), through the mechanism of speciation. For our purpose, species are identified with *genomes*, and genes are arranged linearly or circularly along chromosomes.

#### Reconciled gene trees

A reconciled gene tree is also a binary tree that describes the evolution of a set of genes, called a *gene family*, through the evolutionary mechanisms of speciation, *gene duplication* and *gene loss*, within the given species tree *S*. Therefore, each node of a gene tree *G* represents a gene loss, an extant gene or an ancestral gene. Ancestral genes are represented by the internal nodes of *G*, while gene losses and extant genes are represented by the leaves of *G*.

We denote by *s*(*g*) ∈ *S* the species of a gene *g* ∈ *G*, and by *e*(*g*) the evolutionary event that leads to the creation of the two children *a*_*g*_ and *b*_*g*_. If *g* is an internal node of *G*, then *e*(*g*) is a speciation (denoted by Spec) if the species pair {*s*(*a*_*g*_), *s*(*b*_*g*_)} equals the species pair {*a*_*s*(*g*_), *b*_*s*(*g*_)}, or a gene duplication (GDup) if *s*(*a*_*g*_) = *s*(*b*_*g*_) = *s*(*g*). Finally, if *g* is a leaf, then *e*(*g*) indicates either a gene loss (GLoss) or an extant gene (Extant), in which case *e*(*g*) is not an evolutionary event.

#### Adjacency trees and forests

A *gene adjacency* is a pair of genes that appears consecutively along a chromosome. An adjacency tree represents the evolution of an ancestral adjacency through the evolutionary events of speciation, gene duplication, gene loss (these events, as described above, occur at the gene level and are modelled in the reconciled gene trees), and *adjacency duplication* (ADup), *adjacency loss* (ALoss) and *adjacency break* (ABreak), that are adjacency-specific events.

- The duplication of an adjacency {*g*_1_, *g*_2_} follows from the simultaneous duplication of both its genes *g*_1_ and *g*_2_ (with *s*(*g*_1_) = *s*(*g*_2_) and *e*(*g*_1_) = *e*(*g*_2_) = GDup), resulting in the creation of two distinct adjacencies each belonging to {a_g1_, b_g1_ } × {a_g2_, b_g2_ }.
- An adjacency may disappear due to several events, such as the loss of exactly one (gene loss) or both (adjacency loss) of its genes, or a genome rearrangement that breaks the contiguity between the two genes (adjacency break).

Finally, to model the complement of an adjacency break, i.e. the creation of adjacencies through a genome rearrangement, *adjacency gain* (AGain) events are also considered, and result in the creation of a new adjacency tree. It follows that the evolution of the adjacency between two genes can be described by a forest of adjacency trees, called an *adjacency forest*. In this forest, each node *v* belongs to a species denoted by *s*(*v*), and is associated to an evolutionary event *e*(*v*) ∈ {Spec, GDup, ADup}if *g* is an internal node, or {Extant, GLoss, ALoss, ABreak} if *v* is a leaf. Finally, adjacency gain events are associated to the roots of the trees of the adjacency forest. So in the same way that a gene tree *G* evolves within the species *S*, an adjacency forest *F* describing the evolution of the adjacency between two gene families *G*_1_ and *G*_2_ evolves within *S*, *G*_1_ and *G*_2_. We refer the reader to Fig. 1 for an illustration.

#### Parsimony scores and the Boltzmann distribution

When considered in a parsimonious framework, the score of an adjacency forest *F* is the number of adjacency gains and breaks; other events are not considered as they are the by-products of evolutionary events already accounted for in the score of the reconciled gene trees *G*_1_ and *G*_2_. We denote by *s*_*a*_(*F*) the *parsimony score* of an adjacency forest *F*. Let *F*(*G*_1_, *G*_2_) be the set of all adjacency forests for *G*_1_ and *G*_2_, including both optimal and sub-optimal ones, where we assume that at least one extant adjacency is composed of extant genes from *G*_1_ and *G*_2_.

We define the *Boltzmann factor* of an adjacency forest *F* as

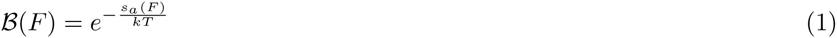

The *partition function* associated to two trees *G*_1_ and *G*_2_ is obtained as

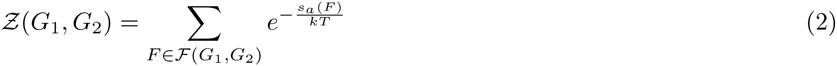

where *kT* is an arbitrary constant. The partition function implicitly defines a *Boltzmann probability distribution* over *Ƒ* (*G*_1_, *G*_2_), where the probability of an adjacency forest *F* is defined by:

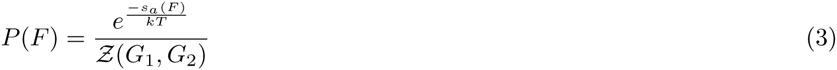

By exponentially favouring adjacency forests with lower parsimony scores, the Boltzmann distribution provides an alternative way to probe the search space, which is heavily influenced by the choice of *kT*. Indeed, decreasing *kT* values will skew the Boltzmann distribution towards more parsimonious adjacency forests. Its limiting distributions are uniform over the whole search space (*kT →* +*∞*) or over the set of co-optimal forests (*kT →* 0) (see Fig. 3 for an illustration).

A Boltzmann probability distribution on the set of all adjacency forests for a given instance also implies a well defined notion of probability for features of adjacency forests. For example, one can associate a probability to a specific potential ancestral adjacency (i.e. adjacency between two genes from a given ancestral species) as the ratio of the sum of the probabilities of the adjacency forests that contain this adjacency with the partition function.

### Algorithms

`DeCo`, the algorithm described in [5] to compute a parsimonious adjacency forest, is a DP scheme constrained by *S*, *G*_1_ and *G*_2_. We first present this algorithm, then describe how to extend it into an Boltzmann ensemble algorithm.

*The DeCo DP scheme.* Let *G*_1_ and *G*_2_ be two reconciled gene trees and *g*_1_ and *g*_2_ be two nodes, respectively of *G*_1_ and *G*_2_, such that *s*(*g*_1_) = *s*(*g*_2_). The `DeCo` algorithm computes, for every such pair of nodes *g*_1_ and *g*_2_, two quantities denoted by *c*_1_(*g*_1_, *g*_2_) and *c*_0_(*g*_1_, *g*_2_), that correspond respectively to the most parsimonious score of a parsimonious adjacency forest for the pairs of subtrees *G*(*g*_1_) and *G*(*g*_2_), under the hypothesis of a presence (*c*_1_) or absence (*c*_0_) of an ancestral adjacency between *g*_1_ and *g*_2_. As usual in DP along a species tree, the score of a parsimonious adjacency forest for *G*_1_ and *G*_2_ is given by min(*c*_1_(*r*_1_, *r*_2_), *c*_0_(*r*_1_, *r*_2_)) where *r*_1_ is the root of *G*_1_ and *r*_2_ the root of *G*_2_.

So, *c*_1_(*g*_1_, *g*_2_) and *c*_0_(*g*_1_, *g*_2_) can be computed as the minimum of a sum of the scores of adjacency gains or breaks and, more importantly, of terms of the form *c*_1_(*x, y*) and *c*_0_(*x, y*) with (x, y) ∈ {g_1_, a_g1_, b_g1_ } × {g_2_, a_g2_, b_g2_ } - (g_1_, g_2_), using the two combinatorial operator min and +.

#### (Un)-ambiguity of the DeCo DP scheme

As defined in [18], the ambiguity of a DP algorithm can be defined as follows: a DP explores a combinatorial solution space (here for `DeCo`, the space of all possible adjacency forests, including possible suboptimal solutions), that can be explicitly generated by replacing in the equations min by ⋒ (the set-theoretic union operator) and + by the Cartesian product *×* between combinatorial sets. A DP algorithm is then unambiguous if the unions are disjoint, i.e. the sets provided as its arguments do not overlap.

We claim that the `DeCo` dynamic programming scheme is unambiguous. Indeed, computing *c*_1_(*g*_1_, *g*_2_) and *c*_0_(*g*_1_, *g*_2_) branches on disjoint subcases that each involve a different set of terms *c*_1_(*x, y*) and *c*_0_(*x, y*). The only case that deserves a closer attention is the case where *e*(*g*_1_) = *e*(*g*_2_) = GDup, as a simultaneous duplication can be obtained by two successive duplications. But in this case, the number of AGain events is different (see Fig. 2), which ensures the pairwise difference of solutions.

**Figure 2.**
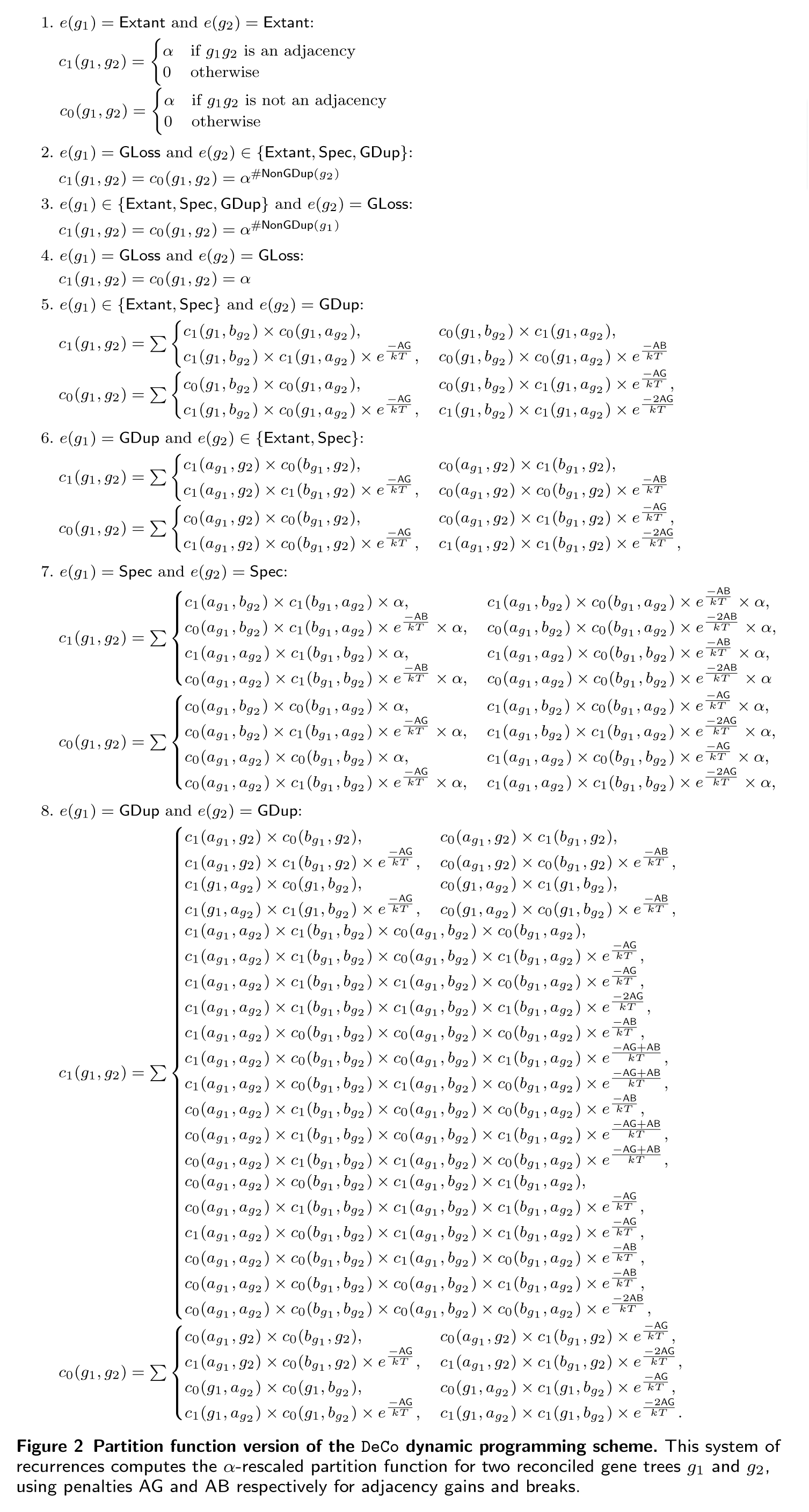
Partition function version of the DeCo dynamic programming scheme. This system of recurrences computes the *α*-rescaled partition function for two reconciled gene trees *g*1 and *g*2, using penalties AG and AB respectively for adjacency gains and breaks.

#### Stochastic backtrack algorithm through algebraic substitutions

As mentioned in [18], any unambiguous dynamic programming scheme can be adapted through algebraic changes to exhaustively generate the set of all adjacency forests, and also compute the corresponding partition function. To that purpose one simply needs), to replace the arithmetic operators (min, +) with (∑, *×*), and to exponentiate any atomic cost *C* ∈ ℝ into a (partial) Boltzmann factor *e*^-*C*/*kT*^ (see Fig. 2).

This precomputation allows us to sample adjacency forests under the Boltzmann distribution, by changing the deterministic backtrack used for maximum parsimony into a stochastic operation. Indeed, assume that the partition function version of the `DeCo` equation computes *c*_1_(*g*_1_, *g*_2_) (resp. *c*_0_(*g*_1_, *g*_2_)) as *i*∈[1,*k*1] ^*t*_*i*_^, where the *t*_*i*_ denote the contribution to the partition function of one of the local alternatives within the DP scheme. The latter are typically computed recursively as combinations of atomic adjacency gain/break costs, and *recursive* terms of the form *c*_1_(*x, y*) and *c*_0_(*x, y*) with (*x, y*) ∈ {*g*_1_, *a*_*g1*_, *b*_*g1*_ } × {*g*_2_, *a*_*g2*_, *b*_*g2*_ } - {(*g*_1_, *g*_2_)}.

Then a (possibly non-parsimonious) random solution can be generated recursively for *c*_1_(*g*_1_, *g*_2_) (resp. *c*_0_(*g*_1_, *g*_2_)), by branching on some *t*_*i*_ with probability *t*_*i*_/*c*_1_(*g*_1_, *g*_2_) (resp. *t*_*i*_/*c*_0_(*g*_1_, *g*_2_)), and proceed recursively on each occurrence of a *recursive* term within the alternative *t*_*i*_. The correctness of the algorithm, i.e. the fact that the random process generates each adjacency forests with Boltzmann probability, follows immediately from general considerations on unambiguous DP schemes [18].

The stochastic nature of the backtrack does not affect its worst-case complexity. This Boltzmann sampling algorithm, for an instance composed of two gene trees *G*_1_ and *G*_2_ of respective sizes (number of leaves) *n*_1_ and *n*_2_, has time complexity of *O*(*n*_1_ *× n*_2_) for each backtrack.

#### Rescaling to avoid numerical overflows

The partition function values *Z*(*g*_1_, *g*_2_), handled during the computation, typically grow exponentially in the total number of nodes in *G*_1_ and *G*_2_, and may end up overflowing the floating point data type used within the DP tables. Following practice in RNA folding prediction [19], we address this issue by iteratively applying an *homogeneous rescaling* of these values during the computation, to keep the values found in the DP table asymptotically close to 1, while still allowing for analysis of the Boltzmann distribution.

To that purpose, one introduces a *rescaling factor α* which is applied, as a multiplicative term, to some of the DP rules. A rescaling is *homogeneous* for a pair of (sub)trees (*G*_1_(*g*_1_), *G*_2_(*g*_2_)) (abridged into (*g*_1_, *g*_2_) from now) when the number of occurrences of *α*, encountered during the generation of a given solution *F*, only depends on (*g*_1_, *g*_2_) and not on specific features of *F*. Let us denote by *κ*_*g1,g2*_ the number of occurrences of *α* for (*g*_1_, *g*_2_), then the rescaled contribution of a given solution *F* is now 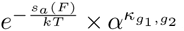, while the *rescaled partition function*, computed by the modified DP scheme, is given by

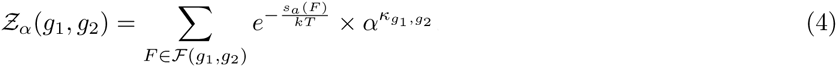

A direct execution of the stochastic backtrack algorithm then returns each forest *F* with probability

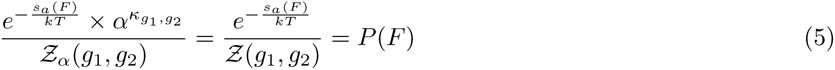

In other words, the introduction of the rescaling does not induce any bias in the stochastic sampling, i.e. the sampling still follows a Boltzmann distribution

**Figure 3.**
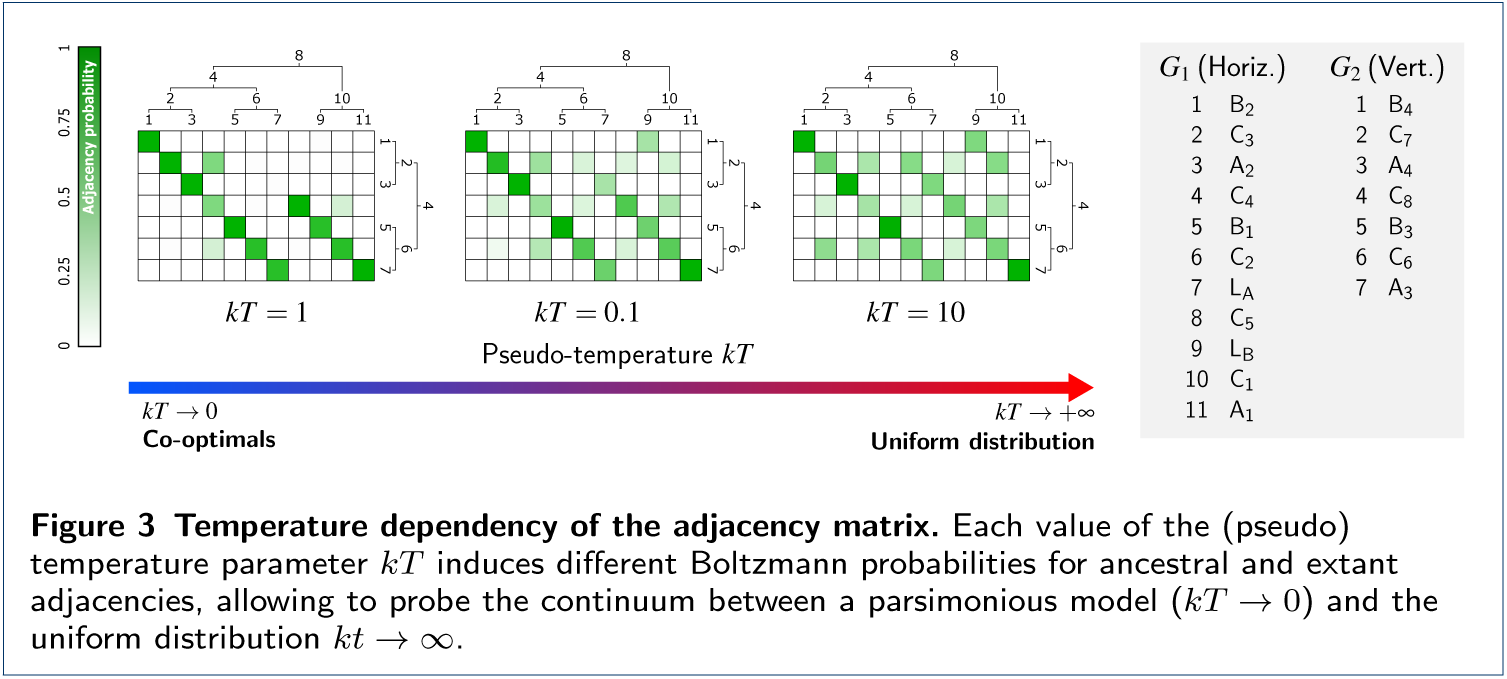
Temperature dependency of the adjacency matrix. Each value of the (pseudo) temperature parameter *kT* induces different Boltzmann probabilities for ancestral and extant adjacencies, allowing to probe the continuum between a parsimonious model (kT → 0) and the uniform distribution kt → ∞.

On the other hand, *α* can be used to constrain the values *Ƶ_α_*(*G*_1_, *G*_2_) to avoid numerical overflows. For instance, setting 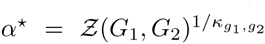 yields *Z*_*α*_*\** (*G*_1_, *G*_2_) = 1. Furthermore, if the rescaling terms are regularly distributed during the execution of the DP scheme, then the intermediate values *c*_0|1_(*g*_1_, *g*_2_) also typically remain close to 1, thereby avoiding numerical over/underflows. In practice, *Z*(*G*_1_, *G*_2_) is the end product of the computation, and thus cannot be used to determine a suitable value for *α*. However, any value that avoids numerical over/underflow can be used, so `DeClone` accepts as input a prescribed value for *α*. Note also that *α* can also be typically inferred from a partial computation, based on the first occurrence of an under/overflow in the DP matrices. To apply these concepts in the context of the `DeCo` DP scheme, we are left to find an homogeneous rescaling.

Fortunately, we observe that the number of recursive calls 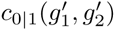, where 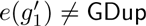 and 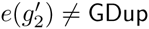, is provably constant^[1]^ within the solutions generated from any call *c*_0|1_(*g*_1_, *g*_2_). From this observation that can be tediously verified by induction, we adapt the DP scheme as illustrated by Fig. 2.

#### Inside-Outside algorithm

While the sampling algorithm described above provides a flexible, easy to implement, approach to analyze the Boltzmann distribution, it only allows for the computation of estimates for properties of interest (for example the occurrence of a specific ancestral adjacency in evolutionary scenarios), whose accuracy may critically depend on the number of samples, the – *a priori* unknown – variance of the underlying distribution, or other factors. However, whenever the property of interest, in conjunction with the DP scheme, fulfills certain technical conditions [18], it is possible to compute its expectation exactly in polynomial time, by transforming the DP scheme using a variant of the *inside-outside algorithm*.

More precisely, our objective is to compute the probabilities associated with each of the *O*(*n*_1_ *× n*_2_) left-hand-side (LHS) to right-hand-side (RHS) transitions in the DP recurrence. Let us denote by *l → r* an LHS/RHS transition, such that

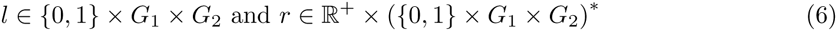

and by *Ƒ*_*l→r*_ the set of forests whose production borrows the *l → r* transition. The Boltzmann probability of (*l → r*) is then defined as

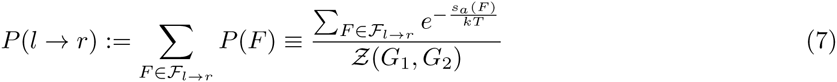

Since *Ƶ* (*G*_1_, *G*_2_) is known, it is sufficient to compute the numerator of the above fraction, i.e. the total Boltzmann factor of the forests *Ƒ*_*l→r*_ that feature (*l → r*). On the other hand, the number of forests in *Ƒ*_*l→r*_ typically grows exponentially on *n*_1_ + *n*_2_, so one must find an efficient strategy for computing this summation.

The principle of the *inside-outside algorithm* [15] is to decompose each of the executions, associated with a forest in *Ƒ*_*l→r*_, into: a) an *inside* part, generated from the recursive calls in the RHS *r*; and b) an *outside* part, which denotes the context in which the LHS *l* appears, i.e. an execution of the DP scheme which features a recursive call to *l*, and is truncated at that point. Let us remark that the inside and outside parts are independent, i. e. any inside part can be combined with any outside part to form a valid execution of the DP scheme, and the score of the associated forest is simply obtained by summing the scores of its two parts. Thus, the total Boltzmann factor of the forests 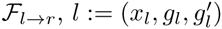, can be decomposed as

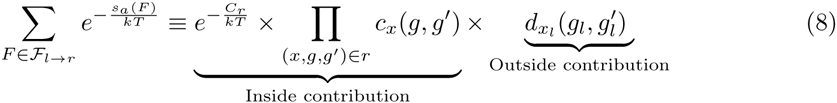

where *C*_*r*_ denotes the constant score increment in the RHS, and 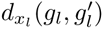 is the outside partition function, i.e. the total Boltzmann factor of all outside parts that are truncated at *l*. This term can be computed in *O*(*n*_1_ *× n*_2_) by *inverting* the DP scheme of Fig. 2 in a purely generic, yet quite technical, fashion [18]. To limit the risk of mistakes in the derivation/implementation of DP equations for *d*_0|1_(*g*_1_, *g*_2_), we implemented an *ad hoc* parser, based on the inversion principle described by Ponty and Saule [18].

Once the probabilities *P* (*l → r*) are known, it is possible to determine the probability of an (ancestral) adjacency (*g*_1_, *g*_2_) by simply summing over the probabilities of transitions that infer such an adjacency, i. e. that feature a recursive call of the form *c*_1_(*g*_1_, *g*_2_) within their RHS. Iterating this over all (*g*_1_, *g*_2_) pairs, one obtains an adjacency matrix, as shown in Fig. 3.

### Results and discussion

#### Data

We re-analyzed a dataset described in [5] composed of 5, 039 reconciled gene trees and 50, 389 extant gene adjacencies, forming 6, 074 DeCo instances, with genes taken from 36 mammalian genomes from the Ensembl database in 2012. In [5], these data were analyzed using the `DeCo` algorithm that computed a single parsimonious adjacency forest per instance. All together, these adjacency forests defined 112, 188 (resp. 96, 482) ancestral and extant genes (resp. adjacencies) ^[2]^, and, more important, lead to 5, 817 ancestral genes participating to three or more ancestral adjacencies, which represent a significant level of syntenic conflict (close to 5%), as a gene can only be adjacent to at most two neighboring genes along a chromosome.

#### DeCo scores, solution space

Unlike reconciled gene trees, whose mutation cost can be high, most adjacency forests have a relatively low cost, with only 32 instances leading to forest of score 5 or above, while the average number of parsimonious syntenic events (adjacency gain and break) is 1.25. This illustrates the fact that syntenic events, that are due to genome rearrangements, are rare evolutionary events, which suggests that parsimony is a relevant criterion for such characters, and that robustness of syntenic characters with respect to the whole solution space should be assessed in terms of optimal or slightly suboptimal evolutionary scenarios.

#### Boltzmann sampling and exact Boltzmann probabilities

For each instance, we sampled 1, 000 adjacency forests under the Boltzmann distribution, for three values of *kT*, 0.001, 0.1, 0.5, and recorded the frequency of all observed ancestral adjacencies. Then for the same values of *kT*, we computed the exact Boltzmann probability of all potential ancestral adjacencies using the inside-outside algorithm. The result observed were very similar whether sampling or exact probabilities were considered. However, the time required to compute exact Boltzmann probabilities is polynomial, so the exact Boltzmann approach based on the inside-outside algorithm should naturally be favoured in applications. In consequence, we discuss only the case of exact Boltzmann probabilities below.

The main difference between the three values of *kT* is that, with *kT* = 0.5, non-optimal adjacency forests have a higher Boltzmann probability in the Boltzmann distribution, while *kT* = 0.1 skews the distribution toward optimal adjacency forests and slightly suboptimal ones, and *kT* = 0.01 ensures that the probability of sub-optimal adjacency forests is extremely low and almost does not contribute to the partition function. We then looked at the numbers of ancestral adjacencies, genes and syntenic conflicts from ancestral adjacencies in terms of Boltzmann probability. Table 1 below summarizes the obtained results.

The difference observed between the results with different values of *kT* supports that parsimony is an appropriate criterion for looking at gene adjacency evolution. Indeed, in the results obtained with *kT* = 0.5, that gives a higher probability to non-optimal adjacency forests, it appears that the number of conserved ancestral adjacencies drops sharply after probability 0.6, showing that very few ancestral adjacencies appear with high probability. However, with *kT* = 0.1 and *kT* = 0.01, by taking a high probability threshold (starting at a threshold of 0.6), we reduce significantly the number of syntenic conflicts while maintaining a relatively similar number of ancestral genes than the experiments described in [5]; this observation illustrates the potential of the ensemble approach compared to the classical dynamic approach that relies on a single arbitrary optimal solution. Next, the experiment with *kT* = 0.01 that considers only co-optimal scenarios (the probability of non-optimal scenarios falls under the numerical precision) shows that, despite conserving only ancestral adjacencies with maximal support in terms of Boltzmann probability, a significant number of syntenic conflicts remains. We conjecture that this is due to errors in the considered reconciled gene trees, and it would be interesting to see if the information about highly supported conflicting adjacencies can be used to correct reconciled gene tree.

**Table 1.**
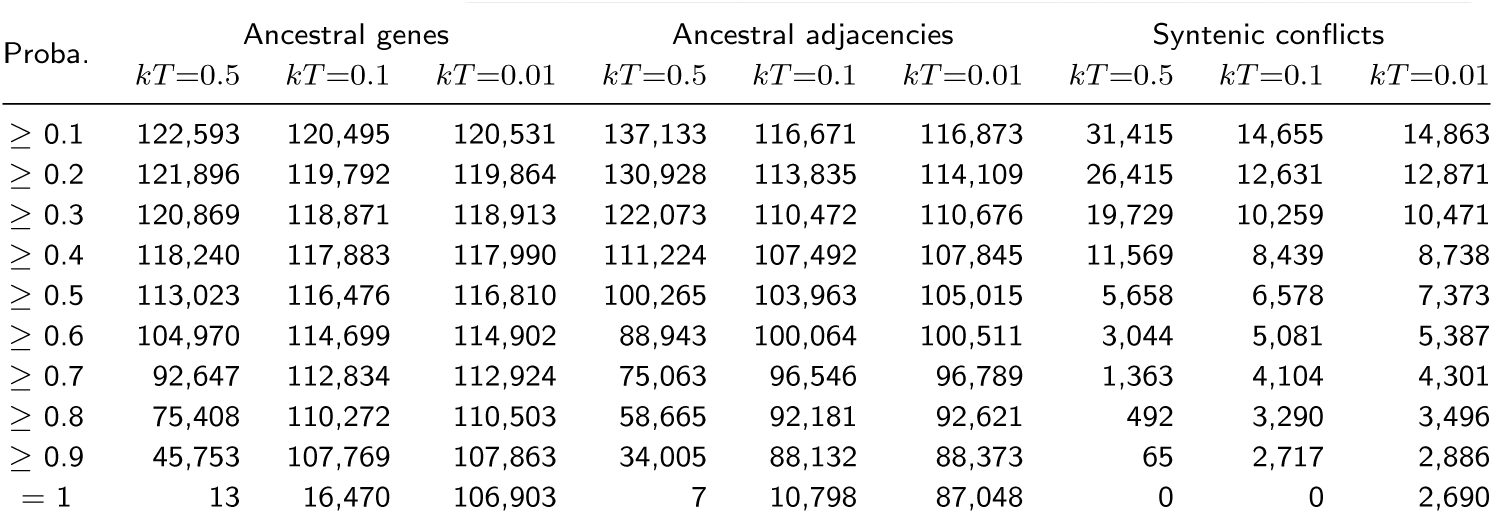
Characteristics of ancestral genes and adjacencies from observed ancestral adjacencies filtered by Boltzmann probability (leftmost column), with different *kT* values.

| Proba. | Ancestral genes |  |  | Ancestral adjacencies |  |  | Syntenic conflicts |  |  |
| --- | --- | --- | --- | --- | --- | --- | --- | --- | --- |
| | $kT=0.5$ | $kT=0.1$ | $kT=0.01$ | $kT=0.5$ | $kT=0.1$ | $kT=0.01$ | $kT=0.5$ | $kT=0.1$ | $kT=0.01$ |
| $\geq 0.1$ | 122,593 | 120,495 | 120,531 | 137,133 | 116,671 | 116,873 | 31,415 | 14,655 | 14,863 |
| $\geq 0.2$ | 121,896 | 119,792 | 119,864 | 130,928 | 113,835 | 114,109 | 26,415 | 12,631 | 12,871 |
| $\geq 0.3$ | 120,869 | 118,871 | 118,913 | 122,073 | 110,472 | 110,676 | 19,729 | 10,259 | 10,471 |
| $\geq 0.4$ | 118,240 | 117,883 | 117,990 | 111,224 | 107,492 | 107,845 | 11,569 | 8,439 | 8,738 |
| $\geq 0.5$ | 113,023 | 116,476 | 116,810 | 100,265 | 103,963 | 105,015 | 5,658 | 6,578 | 7,373 |
| $\geq 0.6$ | 104,970 | 114,699 | 114,902 | 88,943 | 100,064 | 100,511 | 3,044 | 5,081 | 5,387 |
| $\geq 0.7$ | 92,647 | 112,834 | 112,924 | 75,063 | 96,546 | 96,789 | 1,363 | 4,104 | 4,301 |
| $\geq 0.8$ | 75,408 | 110,272 | 110,503 | 58,665 | 92,181 | 92,621 | 492 | 3,290 | 3,496 |
| $\geq 0.9$ | 45,753 | 107,769 | 107,863 | 34,005 | 88,132 | 88,373 | 65 | 2,717 | 2,886 |
| $= 1$ | 13 | 16,470 | 106,903 | 7 | 10,798 | 87,048 | 0 | 0 | 2,690 |

## Conclusions

The main contribution of our work is an extension of the `DeCo` dynamic programming scheme to consider adjacency forests in a probabilistic framework, under the Boltzmann distribution. The application of our algorithms on a mammalian genes dataset, together with a simple threshold-based, approach to filter ancestral adjacencies, proved to be effective to reduce significantly the number of syntenic conflicts, illustrating the interest of the ensemble approach. This preliminary work raises several questions and can be extended along several lines. Among them, we can cite two of immediate interest. First, given the Boltzmann probabilities of the adjacency gains and breaks associated to ancestral adjacencies, we could use them to compute a *Maximum Expected Accuracy* adjacency forest, which is a parsimonious adjacency forest in a scoring model where each event is weighted by Boltzmann probability (see [20] for an example of this approach for RNA secondary structures). This would provide a unique evolutionary scenario per instance. Next, we considered here an evolutionary model based on speciation, duplication and loss. A natural extension would be to include the event of lateral gene transfer in the model. Efficient reconciliation algorithms exist for several variants of this model [3, 4], together with an extension of `DeCo`, called `DeCo`LT [21]. `DeCo`LT is also based on dynamic programming, and it is likely that the techniques we developed in the present work also apply to this algorithm.

## Author’s contributions

All authors participated in all aspects of the study.

## Acknowledgements

J.P.P.Z visit to Simon Fraser University was funded by the São Paulo Research Foundation (FAPESP).

## Footnotes

[1] For the sake of simplicity, we assume here that calls of the form *c*_0|1_(*g*_1_, *g*_2_), where *e*(*g*_1_) = GLoss (resp. *e*(*g*_2_) = GLoss), are expanded into calls *c*_0|1_(*g*_1_, *a*_*g*_2) and *c*_0|1_(*g*_1_, *b*_*g*_2) (resp. *c*_0|1_(*a*_*g*_1, *g*_2_) and *c*_0|1_(*b*_*g*_1, *g*_2_)), unless *g*_2_ (resp. *g*_1_) is also a leaf.

[2] By “ancestral adjacency”, we mean adjacency that involves two genes *g*_1_ and *g*_2_ whose descendants in their respective gene trees satisfy that they do not belong to the same species *s*(*g*_1_) (equal to *s*(*g*_2_)), *i.e. g*_1_ and *g*_2_ are pre-speciation genes, that were not duplicated within their species. This choice is motivated by the fact that the reconstruction of ancestral genomes considers pre-speciation genomes.

## References

1. Liberles, D.A.: Ancestral Sequence Reconstruction. Oxford University Press, Oxford, UK (2007)

2. Csürös, M.: Ancestral reconstruction by asymmetric wagner parsimony over continuous characters and squared parsimony over distributions. In: RECOMB-CG. Lecture Notes in Computer Science, vol. 5267, pp. 72–72. Springer, Berlin, Germany (2008). doi:10.1007/978-3-540-87989-3 6

3. Bansal, M.S., Alm, E.J., Kellis, M.: Efficient algorithms for the reconciliation problem with gene duplication, horizontal transfer and loss. Bioinformatics 28(12), 283–291 (2012). doi:10.1093/bioinformatics/bts225

4. Doyon, J.-P., Scornavacca, C., Gorbunov, K.Y., Szöllosi, G.J., Ranwez, V., Berry, V.: An efficient algorithm for gene/species trees parsimonious reconciliation with losses, duplications and transfers. In: RECOMB-CG. Lecture Notes in Computer Science, vol. 6398, pp. 93–93. Springer, Berlin, Germany (2010). doi:10.1007/978-3-642-16181-0 9

5. Bérard, S., Gallien, C., Boussau, B., Szöllosi, G.J., Daubin, V., Tannier, E.: Evolution of gene neighborhoods within reconciled phylogenies. Bioinformatics 28(18), 382–388 (2012). doi:10.1093/bioinformatics/bts374

6. Biller, P., Feijaö, P., Meidanis, J.: Rearrangement-based phylogeny using the single-cut-or-join operation. IEEE/ACM Trans. Comput. Biology Bioinform. 10(1), 122–134 (2013). doi:10.1109/TCBB.2012.168

7. Gagnon, Y., Blanchette, M., El-Mabrouk, N.: A flexible ancestral genome reconstruction method based on gapped adjacencies. BMC Bioinformatics 13(S-19), 4 (2012). doi:10.1186/1471-2105-13-S19-S4

8. Ma, J., Ratan, A., Raney, B.J., Suh, B.B., Zhang, L., Miller, W., Haussler, D.: DUPCAR: reconstructing contiguous ancestral regions with duplications. Journal of Computational Biology 15(8), 1007–1027 (2008). doi:10.1089/cmb.2008.0069

9. Csürös, M.: How to infer ancestral genome features by parsimony: Dynamic programming over an evolutionary tree. In: Models and Algorithms for Genome Evolution, pp. 29–45. Springer, Berlin, Germany (2013). doi:10.1007/978-1-4471-5298-9 3

10. Libeskind-Hadas, R., Wu, Y.-C., Bansal, M.S., Kellis, M.: Pareto-optimal phylogenetic tree reconciliation. Bioinformatics 30(12), 87–95 (2014). doi:10.1093/bioinformatics/btu289

11. Bansal, M.S., Alm, E.J., Kellis, M.: Reconciliation revisited: Handling multiple optima when reconciling with duplication, transfer, and loss. Journal of Computational Biology 20(10), 738–754 (2013). doi:10.1089/cmb.2013.0073

12. Scornavacca, C., Paprotny, W., Berry, V., Ranwez, V.: Representing a set of reconciliations in a compact way. J. Bioinformatics and Computational Biology 11(2) (2013). doi:10.1142/S0219720012500254

13. Doyon, J.-P., Hamel, S., Chauve, C.: An efficient method for exploring the space of gene tree/species tree reconciliations in a probabilistic framework. IEEE/ACM Trans. Comput. Biology Bioinform. 9(1), 26–39 (2012). doi:10.1109/TCBB.2011.64

14. McCaskill, J.S.: The equilibrium partition function and base pair binding probabilities for RNA secondary structure. Biopolymers 29(6-7), 1105–1119 (1990). doi:10.1002/bip.360290621

15. Baker, J.K.: Trainable grammars for speech recognition. The Journal of the Acoustical Society of America 65(S1), 132–132 (1979). doi:10.1121/1.2017061

16. Arvestad, L., Lagergren, J., Sennblad, B.: The gene evolution model and computing its associated probabilities. J. ACM 56(2) (2009). doi:10.1145/1502793.1502796

17. Mahmudi, O., Sjöstrand, J., Sennblad, B., Lagergren, J.: Genome-wide probabilistic reconciliation analysis across vertebrates. BMC Bioinformatics 14(S-15), 10 (2013). doi:10.1186/1471-2105-14-S15-S10

18. Ponty, Y., Saule, C.: A combinatorial framework for designing (pseudoknotted) RNA algorithms. In: Przytycka, T., Sagot, M.-F. (eds.) Algorithms in Bioinformatics (Proceedings of WABI’11). Lecture Notes in Computer Science, vol. 6833, pp. 250–250. Springer, Berlin Heidelberg, Germany (2011). doi:10.1007/978-3-642-23038-7 22

19. Hofacker, I.L., Fontana, W., Stadler, P.F., Bonhoeffer, L.S., Tacker, M., Schuster, P.: Fast folding and comparison of RNA secondary structures. Monatshefte für Chemie / Chemical Monthly 125(2), 167–188 (1994). doi:10.1007/BF00818163

20. Clote, P., Lou, F., Lorenz, W.A.: Maximum expected accuracy structural neighbors of an RNA secondary structure. BMC Bioinformatics 13(S-5), 6 (2012). doi:10.1186/1471-2105-13-S5-S6

21. Patterson, M., Szöllosi, G.J., Daubin, V., Tannier, E.: Lateral gene transfer, rearrangement, reconciliation. BMC Bioinformatics 14(S-15), 4 (2013). doi:10.1186/1471-2105-14-S15-S4

